# Multi-scale neural networks enhance species distribution modelling across predictors and taxonomic groups

**DOI:** 10.1101/2025.10.15.682577

**Authors:** Nina van Tiel, Robin Zbinden, Chiara Vanalli, Benjamin Kellenberger, Loic Pellissier, Devis Tuia

## Abstract

Ecological processes and patterns are scale-dependent, as reflected in how relationships between species occurrence and the environment vary across scales. In species distribution models (SDMs), spatial scale can strongly affect model predictions and performance. Despite this, the choice of scale is often overlooked. The development of SDMs using deep learning models has enabled spatially structured data, such as patches of environmental raster data, to be considered as model inputs, making the question of scale even more crucial. Here, we evaluate convolutional neural network-based multi-species SDMs considering different scales on a dataset from Switzerland, including more than 8 million observations for 2390 plant and 1006 animal species comprising amphibians, butterflies, beetles, and mammals. We investigate how scale affects model performance and compare single- and multi-scale models. Our results reveal stronger scale effects for remote sensing indices than for bioclimatic and edaphic variables. In single-scale models, we find that plant species perform better with smaller spatial scales, while the opposite is true for animal species. Multi-scale models consistently improve predictive performance and reduce sensitivity to arbitrary scale selection across all modalities and taxonomic groups. The effect of scale is nonetheless smaller than the increase in performance achieved by considering multiple predictor groups. To improve the interpretability of such complex models, we use attribution methods to compute the relative contribution of different scales and predictor groups. Our study demonstrates the importance of accounting for scale and the potential of deep learning methods to integrate ecological complexity with spatial data across scales.

**Highlights:**

- Deep learning SDMs can use spatial environmental data as model predictors
- The spatial extent of the predictors affects model performance
- Multi-scale models improve performance across predictors and taxonomic groups
- Multi-scale modelling reduces sensitivity to arbitrary scale selection
- Explainable AI methods provide contribution scores for scales and predictor groups

## 1. Introduction

Understanding the geographic distribution of species is a prerequisite for mapping biodiversity and guiding the protection and restoration of nature. A prevalent approach to infer the distribution of species is to relate species occurrence records to environmental variables (Guisan and Thuiller, 2005). Species distribution models (SDMs) are used to gain a better understanding of biogeographic patterns or to predict species distributions across a landscape (Elith and Leathwick, 2009), and are invaluable tools to support decision-making for conservation (Guisan et al., 2013). Since ecological processes that shape species distributions operate at different scales (Guisan and Thuiller, 2005; McGill, 2010), the relationship between species occurrence and the environment can depict different spatial patterns at different scales (Wiens, 1989; McGarigal et al., 2016; Lu and Jetz, 2023). For example, climatic variables are considered to shape large-scale distribution patterns, whereas biotic effects or habitat may be more important at local scales (König et al., 2021; Wisz et al., 2013). Accordingly, the scale at which environmental variables are considered has a strong impact on the inferred species-environment relationships, impacting the performance of SDMs (Chauvier et al., 2022). Furthermore, different taxonomic groups of species have different ecological requirements and drivers operating at different scales in relation to their biological traits (Turner et al., 2019; Manzoor et al., 2018; Guisan et al., 2007).

Over the past decades, many statistical models have been proposed to model species’ distributions, with correlative, occurrence-based approaches being the most widely used in the field (Elith et al., 2010). Modelling choices, ranging from the type of data to the algorithms used, have strong implications on the suitability and appropriateness of the output for diverse applications and objectives (Guillera-Arroita et al., 2015). This includes the choice of geographic scale of the analysis, including both the spatial extent and the resolution of the predictor. Scale can affect not only a model’s predictive performance but also the conclusions drawn from downstream analyses of the resulting distribution maps (Chauvier et al., 2022), for example, when using SDMs for climate change impact assessments (Nadeau et al., 2017). A study considering high-elevation plant species in two regions of Switzerland illustrates this issue (Randin et al., 2009): a model using coarse climatic data (10′ × 10′ grid cells) predicted that 52% of species would become extirpated under climate change, while 100% of species were predicted to persist when considering high-resolution data (25m × 25m grid cells). Nonetheless, scale dependency is often overlooked in species distribution modelling. A recent review found that the spatial grain used for environmental niche analysis ranged from 1 m to 300 km (Lu and Jetz, 2023). They found that the grain size varied significantly among taxonomic groups, with smaller scales used for less mobile species, such as plants and invertebrates. Generally, the scale roughly matched the sampling resolution of the readily available biodiversity and environmental data, indicating that the choice of grain size may rely predominantly on the scale of widely available environmental data (≈1 km) rather than prior knowledge about the appropriate scale (Lu and Jetz, 2023; Manzoor et al., 2018).

The vast majority of SDM approaches, such as MaxEnt (Phillips and Dudík, 2008), generalised linear models (Chiaverini et al., 2023), or random forest models (van Tiel et al., 2024), as well as more recent deep learning-based models (Hu et al., 2025), consider tabular data of environmental variables, with each predictor represented by a single value for a given location (Valavi et al., 2022). Therefore, the scale of the predictors associated with each observation is inherently linked to the grain size of the underlying data. Higher resolution spatial data, such as satellite images or microclimatic data derived from remote sensing, may present new opportunities to explore finer-scale patterns (Nadeau et al., 2017; Randin et al., 2020). However, models that handle tabular data are limited by their inability to capture spatial patterns in environmental variables.

In contrast, neural networks, or deep learning models in general, provide a striking flexibility in the number and dimensionality of inputs, allowing the inclusion of non-tabular data. In particular, convolutional neural networks (CNNs) have been proposed for species distribution modelling to overcome this limitation (Deneu et al., 2021). CNNs are a type of artificial neural network that can capture spatial patterns in images and have been successfully applied in computer vision (Li et al., 2021), including ecological applications (Brodrick et al., 2019; Tuia et al., 2022; Pollock et al., 2025). When used as SDMs, the spatial context around observations, beyond the granularity of the predictors, can be considered by the model (Botella et al., 2018). In this context, scale is related to two factors: the grain size of the data and the spatial extent of the image patches. We will use the term “resolution” to refer to the former and “scale” to refer to the latter. SDMs have been constructed with CNNs applied to patches extracted from rasters of environmental variables (Botella et al., 2018; Deneu et al., 2021), satellite images (Estopinan et al., 2022; Dollinger et al., 2024; Hu et al., 2025), and both together (Teng et al., 2024; van Tiel et al., 2025). Although the inclusion of spatial data describing the environment can lead to higher numeric model performance (Deneu et al., 2021), the optimal scale at which these predictors are relevant to the model remains an open question. While the resolution of spatial data varies, the size of the image patches considered by these models is rarely justified, and their corresponding spatial extent varies extensively, ranging from 0.01 to 4000 km^2^ in the studies cited above.

Moreover, neural networks enable modelling the distribution of many species with a single model. As the appropriate scale may differ among species (Guisan et al., 2007), the selection of the “best” scale for a model is not evident, and the imposition of a single scale for all species is debatable (Wiens, 1989; Lu and Jetz, 2023). Similarly, imposing a single scale for all predictors is also questionable (Fournier et al., 2017). The idea of adopting a multi-scale perspective in ecological models to address this matter is far from novel (Wiens, 1989; Hallman and Robinson, 2020), although it is not standard in species distribution modelling. The flexibility of deep learning models can facilitate overcoming the limitation of fixed scales. For instance, multi-scale models from computer vision (Ronneberger et al., 2015; Chen et al., 2017) can be adapted to SDMs, allowing the context around a species observation to be simultaneously considered at different extents. The flexible design of deep learning models also enables the construction of architectures that can take into account diverse types of predictors, or modalities. Such multimodal models have been developed in the context of SDMs to combine, for instance, point values of environmental predictors with satellite imagery (Dollinger et al., 2024; Hu et al., 2025). Spatial data considered at different scales or resolutions may also be treated as modalities. For example, bioclimatic variables are typically available at around 1 km and satellite imagery at much finer resolutions (McGill, 2010). To enable variability in the scales for each modality, a late fusion strategy, where information streams from different modalities are merged in the last layer of the neural network, allows each modality to be considered at its native resolution and different scales.

A recent study presented a CNN-based multimodal multi-scale SDM using rasters of bioclimatic variables and satellite images with plant occurrence data across Europe (van Tiel et al., 2025). Their modelling framework addresses the limitations stated above. Their results indicate that the scale of both environmental variables and satellite images affects model performance, with multi-scale strategies benefiting satellite images more than the bioclimatic predictors. However, with the inclusion of several scales and modalities, it is pertinent to try to gain a better understanding of which factors contribute most to the final predictions. Deep learning models are generally considered opaque, black boxes, but tools from explainable artificial intelligence (xAI) have provided us with methods to better understand their decision process (Arrieta et al., 2020). For instance, in computer vision, attribution methods generate visual explanations that highlight which part of the image influences the model output (Ancona et al., 2019). xAI methods have only been used sparingly in SDMs so far, yet they can provide valuable insights about which predictors are most important for the model’s decision process (Bourhis et al., 2023; Hu et al., 2025; Zbinden et al., 2025).

In this study, we investigate scale effects in a multimodal and multi-scale SDM framework, based on van Tiel et al. (2025). We extend the taxonomic scope by considering vascular plants, as well as amphibian, butterfly, beetle, and mammalian species across Switzerland, to evaluate how scale dependencies vary across the tree of life. Species occurrence data are related to patches of very high-resolution (25 m) environmental predictors from `SWECO25` (Külling et al., 2024) of varying spatial extents (125 – 3375 m). We investigate how spatial scale affects model performance and whether multi-scale approaches outperform single-scale models. We further evaluate the effect of including different environmental data modalities, namely bioclimatic, edaphic, and remote sensing-derived variables, both in uni- and multimodal model configurations. We also examine how these effects may vary across taxonomic groups. Finally, we use a gradient-based xAI method to quantify how different scales and groups of predictors contribute to multi-scale and multimodal model outputs across species and landscapes. We find that relationships between scale and model performance differ across modalities, with the strongest scale effects found for remote sensing indices. Optimal single scales vary across taxonomic groups, with plants favouring smaller scales than animals. Moreover, our results indicate that multi-scale and multimodal approaches consistently lead to better model performance for all taxonomic groups considered here. Overall, this study underlines the importance of scale and the capability of deep learning methods to use spatial data at multiple scales, thus addressing the scale dependence of species-environment relationships.

## 2. Materials and methods

### 2.1. Occurrence data

The species occurrences used in this study consist of opportunistic observations from citizen science for plant and animal species in Switzerland collected between 2000 and 2025. The National Data and Information Center on the Swiss Flora (InfoFlora, 2025) provided data for vascular plant species. Data for amphibians, terrestrial mammals, beetles, and butterflies were obtained from the National Data and Information Center on the Swiss Fauna (InfoFauna, 2025). Together, 11, 612, 211 occurrence records were obtained for 4232 plant species and 1569 animal species. For plant data, we retained only wild-growing or officially reintroduced, indigenous or archaeophytic plant occurrences, according to the InfoFlora database, removing 14% of all occurrences. For animals, we removed non-native species, according to the InfoFauna database, and observations with a location uncertainty larger than 1 km, together accounting for 1.2% of occurrences. No location uncertainty filter was applied to the plant occurrences, as the maximum uncertainty was 100 m. The coordinates were transformed to the same coordinate reference system as the predictors (`EPSG:2056`), and duplicate observations of the same species in the same 25 × 25 m grid cell were removed. To avoid overoptimistic model evaluation due to spatial autocorrelation associated with random data splitting (Roberts et al., 2017), we use spatial blocks of 25 × 25 km to split the occurrence data into train, validation, and test sets, such that the number of locations per split was approximately 70*/*15*/*15% for training, validation, and test sets, respectively (Figure 1a and S1). Finally, we retain the occurrences of taxa with at least 10 occurrences in the training data and at least one occurrence in the validation and test splits. Altogether, we are left with 8, 456, 749 occurrences spread over 2, 426, 551 locations for 3396 taxa: 18 Amphibia, 547 Coleoptera, 383 Lepidoptera, 58 Mammalia and 2390 Plantae. The number of occurrences per species in the training data shows a long-tailed distribution (Figures 1b and S1).

**Figure 1:**
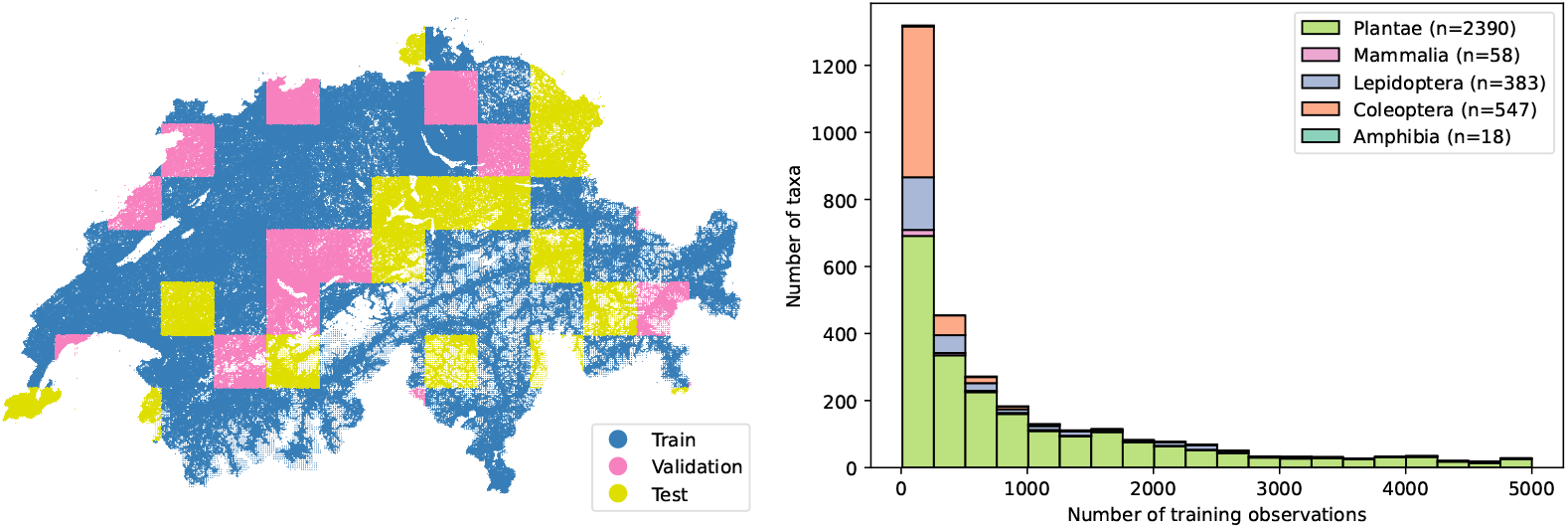
Occurrence data distribution. (a) Geographic distribution of training, validation, and test splits with spatial blocks. (b) Stacked histogram of the number of occurrences per species in the training set, coloured by taxonomic group. Separate plots for each taxonomic group can be found in Figure S1.

### 2.2. Environmental predictors

We use 39 high-resolution (25 m) rasters from the `SWECO25` dataset as model predictors (Külling et al., 2024). We consider three groups of variables. The first modality is bioclimatic variables, containing 19 layers representing annual trends, seasonality, and extremes, averaged over 30 years (1981–2010). These layers were downscaled from 1 km to 25 m using local regressions with an elevation model to account for local topography. The second modality contains 10 layers describing soil properties and were resampled from their original spatial resolution of approximately 100 m to 25 m. The third modality consists of remote sensing indices, which were resampled from their original resolution of 10 m. Yearly mean and standard deviation layers for five vegetation indices were available and we selected the 10 layers from the year 2018, as it was the most recent year for which all indices were available. Before training our models, each raster layer is standardised by subtracting its mean and dividing by its standard deviation.

### 2.3. Model architecture

We construct our models based on the model architecture presented in (van Tiel et al., 2025), which allows us to construct models that consider one or several modalities, at one or multiple scales. We modify the exact sequence of convolutions, including the number, size, and arrangement of convolutional and pooling layers, to accommodate our 25 m resolution predictor variables, rather than the 1 km and 10 m resolutions for bioclimatic variables and remote sensing images, respectively, used in the original study. The model, shown in Figure 2, is structured into three parts. First, the *encoder* consists of two convolutional layers with 64 filters of size 3 × 3 each, resulting in a receptive field of 5 × 5 pixels. Batch normalisation and ReLU are applied after each convolutional layer. Second, the *spatial module* has one or multiple branches for single- or multi-scale models. Each branch consists of repetitions of a spatial block, each consisting of two 3 × 3 convolutional layers with 64 filters, separated by a 3 × 3 max pooling operation. The central pixel of the resulting tensor is then extracted to generate a 64-dimensional vector. The receptive field of the central pixel corresponds to the scale considered by that branch of the spatial module. Finally, the output vectors of the spatial modules, each corresponding to a given scale and modality, are concatenated before the final *classification layer* in a late fusion approach (Figure 2d, e). The final layer is a linear layer with 3396 outputs, corresponding to the number of species. The sigmoid function is applied to the output vector to obtain predictions between 0 and 1. This enables the model to predict the suitability of a site for each species independently, allowing multiple species to be predicted as present at each site, in a multi-label classification fashion (Hu et al., 2025).

**Figure 2:**
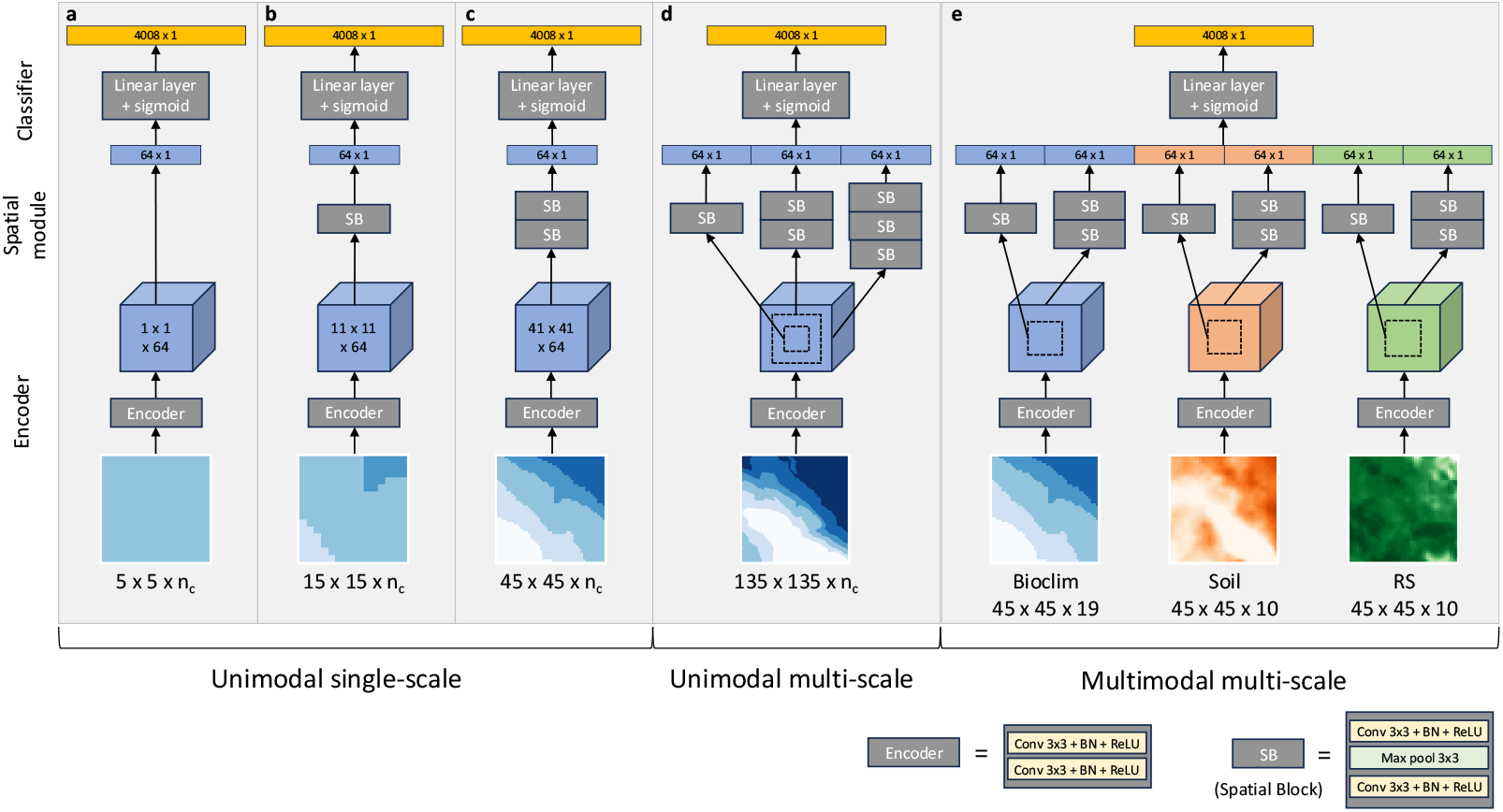
Examples of model architectures with the modular structure for SDMs. (a-c) Single-scale, unimodal model architecture for bioclimatic variables at scale 5 × 5, 15 × 15, and 45×45 pixels. (d) Multi-scale unimodal model architecture for bioclimatic variables at scales 15 × 15, 45 × 45, and 135 × 135 pixels. (e) Multi-scale multimodal model architecture for bioclimatic variables, soil variables, and remote sensing indices (RS) at scales 15 × 15 and 45 × 45 pixels.

To investigate the role of the scale of predictors, we trained models considering spatial extents varying from 0.02 km^2^ to 11.39 km^2^. While this framework allows different modalities to be considered at different scales, all our predictors have the same resolution of 25 × 25m. We therefore use the same model architectures for all modalities, and the same scales are tested for each group of predictors. Specifically, we construct single-scale models applying the spatial block between 0 and 3 times, leading to scales of 5 × 5, 15 × 15, 45 × 45, and 135 × 135 pixels, or 125 × 125, 375 × 375, 1125 × 1125 and 3375 × 3375 m (Figure 2a-c). As all patches are square, we will hereafter refer to each scale by its side length (i.e., 5, 15, 45, and 135). Multi-scale models that consider two or three scales are also tested (e.g., Figure 2d). We evaluate the effect of scale for each of the three modalities, bioclimatic, soil, and remote sensing indices, separately in unimodal models (Figure 2a-d), as well as together in multimodal models (Figure 2e).

### 2.4. Model training and evaluation

We train our models with a weighted binary cross-entropy loss (Zbinden et al., 2024). This loss function is adapted for multi-label classification, allowing multiple species to be predicted at each site by comparing the model predictions to the ground truth with species-specific weights. These weights aim to address the class imbalance, in which there are drastically different numbers of occurrences for the different taxa in the training data (Figure 1b). Here, we consider the records of other species as pseudo-absences. Model parameters are learned with a schedule-free stochastic gradient descent optimiser (Defazio et al., 2024) with a learning rate of 0.1 and a batch size of 64. We train each model for 30 epochs. We evaluate models with the area under the receiver operating curve (AUC) and compute the median AUC across all species, as well as for the various taxonomic groups. To avoid overfitting on the training data, we retain the model weights from the epoch that yields the best median AUC on the validation dataset for model testing.

### 2.5. Gradient-based model explanation

To better understand which scales and modalities are most important for model predictions, we use a gradient-based attribution method in order to estimate the contribution of the features associated with each scale and modality. Gradients, most commonly first-order derivatives, are used to train deep learning models: computed for each model parameter with respect to the value of the loss function, they indicate the direction and magnitude by which the parameters need to be adjusted to minimise the loss value. In gradient-based explainability methods, gradients may instead be computed for input variables with respect to the model predictions themselves. In this case, they provide a local estimate of the direction and strength in which each input has to change to maximise the predicted output for a given sample. However, raw gradients are generally not considered a trustworthy local approximation because of the discontinuous nature of the model’s prediction function (Shrikumar et al., 2017). Multiplying the gradient with the input values leads to sharper attribution patterns and has therefore been proposed as a better measure of the contribution of the input variables to the model output (Simonyan et al., 2013; Ancona et al., 2017). Here, we consider the feature space before the final classification layer of our neural network, which consists of concatenated 64-dimensional vectors, each corresponding to one scale and modality. Guided by the goal of integrating interpretability, the late-fusion design of our models separates data streams by scale and modality, enabling gradient-based attribution methods to operate directly in the feature space. We quantify the contribution of each element of the feature space to the final prediction for a given sample by multiplying each feature by its gradient with respect to the model prediction for a given species. The contribution of a given modality and scale is the sum of the absolute values of the gradient × feature of the 64 corresponding elements of the feature vector. We normalise them such that the sum of contributions for each species is equal to 1. We calculate local explanations in the form of relative contributions for each modality and scale in a given model at 5000 locations sampled along a grid that covers Switzerland. To obtain a global explanation, we consider the mean relative contribution across these samples for each species.

## 3. Results

### 3.1. Scale effects across modalities

We investigate the role of the scale of predictors by comparing the performance of models on a spatial-block held-out test set for scales of 5, 15, 45, and 135 pixels. Examining the median performance across all species (Figure 3a, Table S1), we find that scale affects model performance with a variation in intensity and effect across modalities. Our results indicate that the effect of scale is strongest on remote sensing indices, with a monotonically increasing relationship to predictive performance in single-scale settings. Using the 5-pixel scale yields the lowest median AUC scores of all tested models (78.4%). However, model performance increases to values comparable to or surpassing those of models using bioclimatic variables for larger scales and multi-scale settings (81.2 − 83.9%). Edaphic factors have the best performance among unimodal models across all scales (85.8 − 88.5%). The relationship between model performance and scale is less pronounced for bioclimatic and soil variables, with a median coefficient of variation (COV) of model performance across single-scale models of 1.9% and 1.7%, respectively, compared to 3.0% for remote sensing indices. For these two modalities, the best single-scale model performance is obtained with intermediate scales of 15 or 45 pixels, and drops in performance are observed for the smallest and largest spatial extents. Multimodal models, which use all three predictor groups, depict the same trend with less variability (median COV of 1.5% across single-scale models) and achieve the best performance for all scales (87.3 − 89.2%).

**Figure 3:**
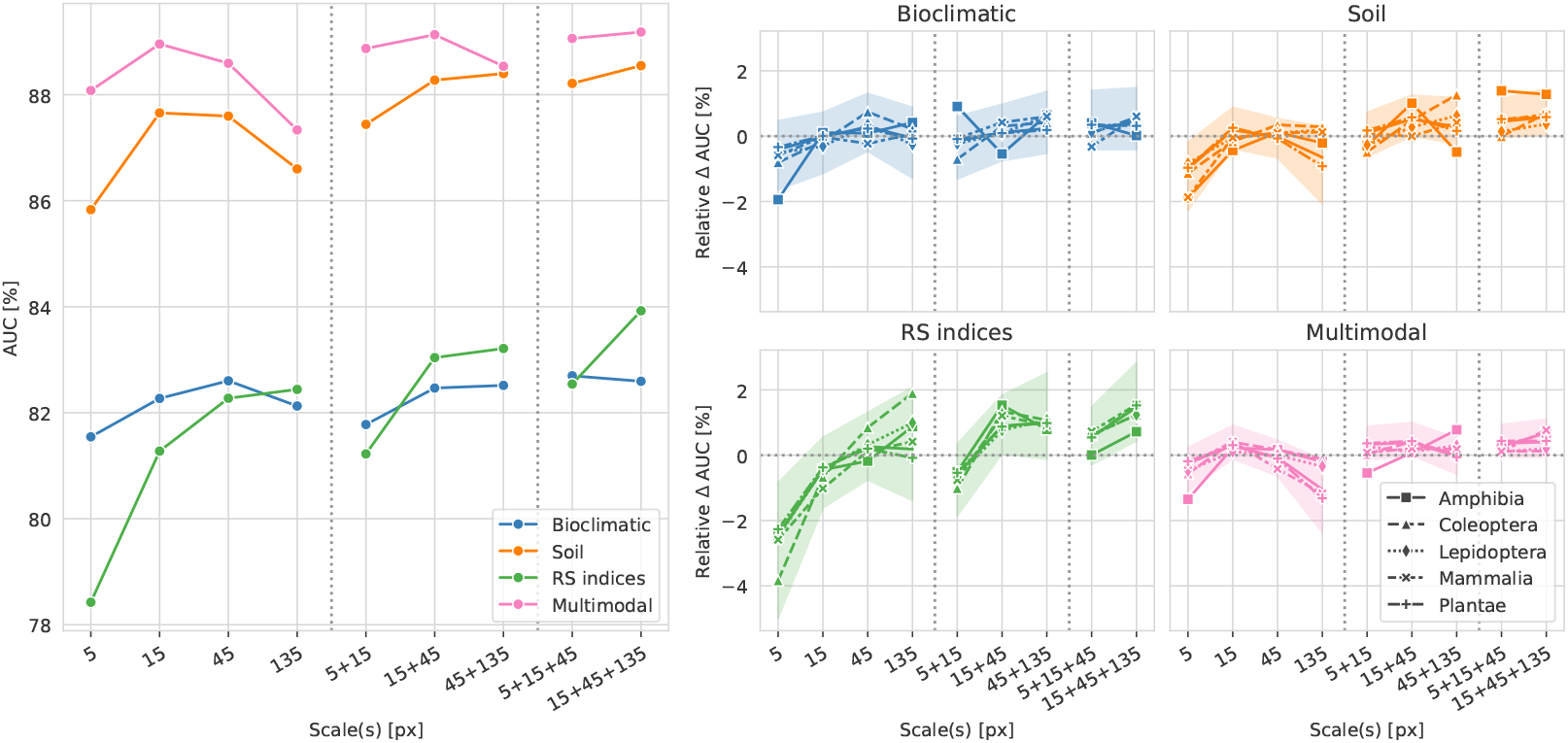
(a) Median AUC scores (%) evaluated on the test set for different scales for bioclimatic variables, soil variables, remote sensing (RS) indices, and all of the above (Multimodal). Dotted lines separate single scales, scale pairs, and triplets. (b) Relative difference in AUC (%) computed for each species and scale as the difference between the scale-specific AUC and the mean AUC across scales for each modality. Positive values indicate higher performance than the mean AUC and vice versa. Markers indicate the median for each group of species, and the shaded area is the interquartile range across all species.

### 3.2. Single- vs. multi-scale models

We also evaluate models trained on multiple scales by considering pairs and triplets of consecutive scales. We find that multi-scale models generally match or exceed the median AUC of their single-scale counterparts, with models with triplets of scales leading to the best median AUC for each modality (82.7%, 88.5%, 83.9%, 89.2% for bioclimatic, edaphic, remote sensing variables, and all of the above, respectively) (Figure 3a, Table S1). The increase in predictive performance compared to single-scale models is, nonetheless, limited. For bioclimatic variables, the median gain between median AUC scores for multi-scale and single-scale models is +0.21%. The gain is most evident for remote sensing variables (+1.20%). Moreover, multi-scale models lead to more stable model performance with lower variability in AUC than single-scale models. Models trained on remote sensing indices are subject to the largest decrease in variability, with a median difference in COV between multi- and single-scale models of −0.8%, although they retain the largest median COV of all modalities of 2.1%. The median difference in COV is marginal for models using bioclimatic variables (−0.1%). Both edaphic-based and multimodal models have a median decrease of COV of −0.4%, resulting in the lowest multi-scale COVs of 1.0% and 0.9%, respectively.

### 3.3. Scale effects across taxonomic groups

To compare scale effects across taxonomic groups, we investigate the relative effect of scale by computing the difference between scale-specific AUC and the mean AUC across scales (relative ΔAUC) for a given modality and for each species. We find the shape and relative magnitude of scale effects are generally consistent across taxonomic groups for each modality (Figure 3b). These results reflect the strong scale effects for remote sensing indices compared to other modalities, in particular for Coleoptera taxa. The Amphibia class also stands out with a large relative ΔAUC for bioclimatic and soil variables for some scales. However, this may be due to the small number of species in this group.

Absolute model performance per taxonomic group (Figure S2 and Table S1) reveals that smaller scales of 15 and 45 pixels lead to the best single-scale median AUC for plant species. In comparison, animal taxonomic groups reach their best single-scale performance most often with the largest scale of 135 pixels. Multi-scale models outperform single-scale models for all taxonomic groups for two of the three modalities, except Lepidoptera, for which they were superior only for soil variables. When all three modalities are considered, multi-scale models are superior across all taxonomic groups. Multi-scale approaches seem to be particularly beneficial for amphibian taxa with relatively large median gains in predictive performance between the median AUC scores for multi-scale and single-scale models (+0.5%, +1.3%, +1.0%, +0.6% for bioclimatic, soil, remote sensing indices, and multimodal, respectively). In contrast, our results indicate that Coleoptera have the smallest gains from multi-scale modelling (+0.1%, +0.5%, +0.9%, +0.1%). Furthermore, we find strong variation in absolute model performance across taxonomic groups, which may surpass the variation in AUC scores within a group across different scales or modalities (Table S1 and Figure S2). Mammalia taxa obtain the lowest model performances (71.0 − 77.9%), while Plantae obtain the highest median AUC scores (80.3 − 90.7%). Amphibia, Coleoptera, and Lepidoptera taxa have intermediate median AUC scores. The ranking of modalities is consistent across taxonomic groups.

### 3.4. Relative contributions of scales and modalities

We further investigate the relative contribution of each modality and scale for one of the best-performing models: the multimodal model considering scales of 15 and 45 pixels. We compute the relative contributions for 5000 sites sampled along a regular grid. When aggregated across all sites, we find that soil variables on a scale of 15 pixels are considered the most important by the model for all taxa (Figure 4a). Soil variables on a scale of 45 pixels and bioclimatic variables and remote sensing indices on a scale of 15 pixels follow. Finally, bioclimatic variables and remote sensing indices on a scale of 45 pixels are the least important predictors according to our explainability metrics. These results, in particular the strong contribution of edaphic factors, are consistent with our results on the performance of unimodal models. Our results indicate that the ranking of the relative contribution of the various modalities and scales is consistent across taxonomic groups (Figure 4b). We find that amphibian species show the most variability compared to the other taxa.

**Figure 4:**
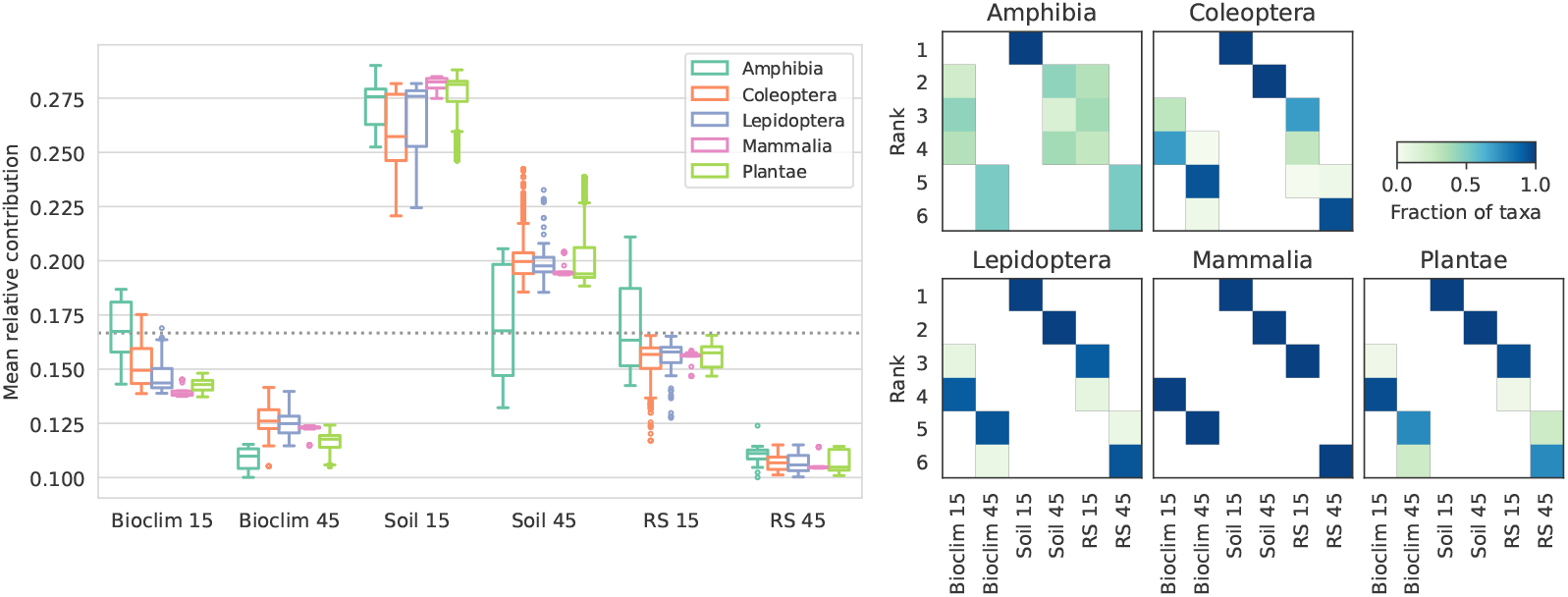
(a) Boxplots of mean relative contribution of modalities and scales for a multimodal multi-scale model (scales 15 and 45) computed per taxonomic group for 5000 samples randomly sampled from the test set. (b) Heatmaps of the rank of mean relative contribution of modalities and scales for each taxonomic group.

Moreover, we consider the local relative contributions of the different modalities for specific species to illustrate spatial differences in variable importance. Here, we present maps of relative modality importance, where contributions for the two scales of each modality are summed and mapped to cyan, yellow, and magenta for bioclimatic, soil, and remote sensing variables, respectively, for one species selected for each taxonomic group (Figure 5). While the geographic distributions of species differ, the patterns of the relative contribution of modalities are similar for four of the five species shown here. Soil variables have the largest contribution to model predictions overall, but they are complemented by the contributions of bioclimatic factors in the Alps and remote sensing indices in the Swiss Plateau. The results for the European tree frog, as an example species for amphibians, differ from these patterns with a strong contribution of remote sensing indices across a large part of the landscape.

**Figure 5:**
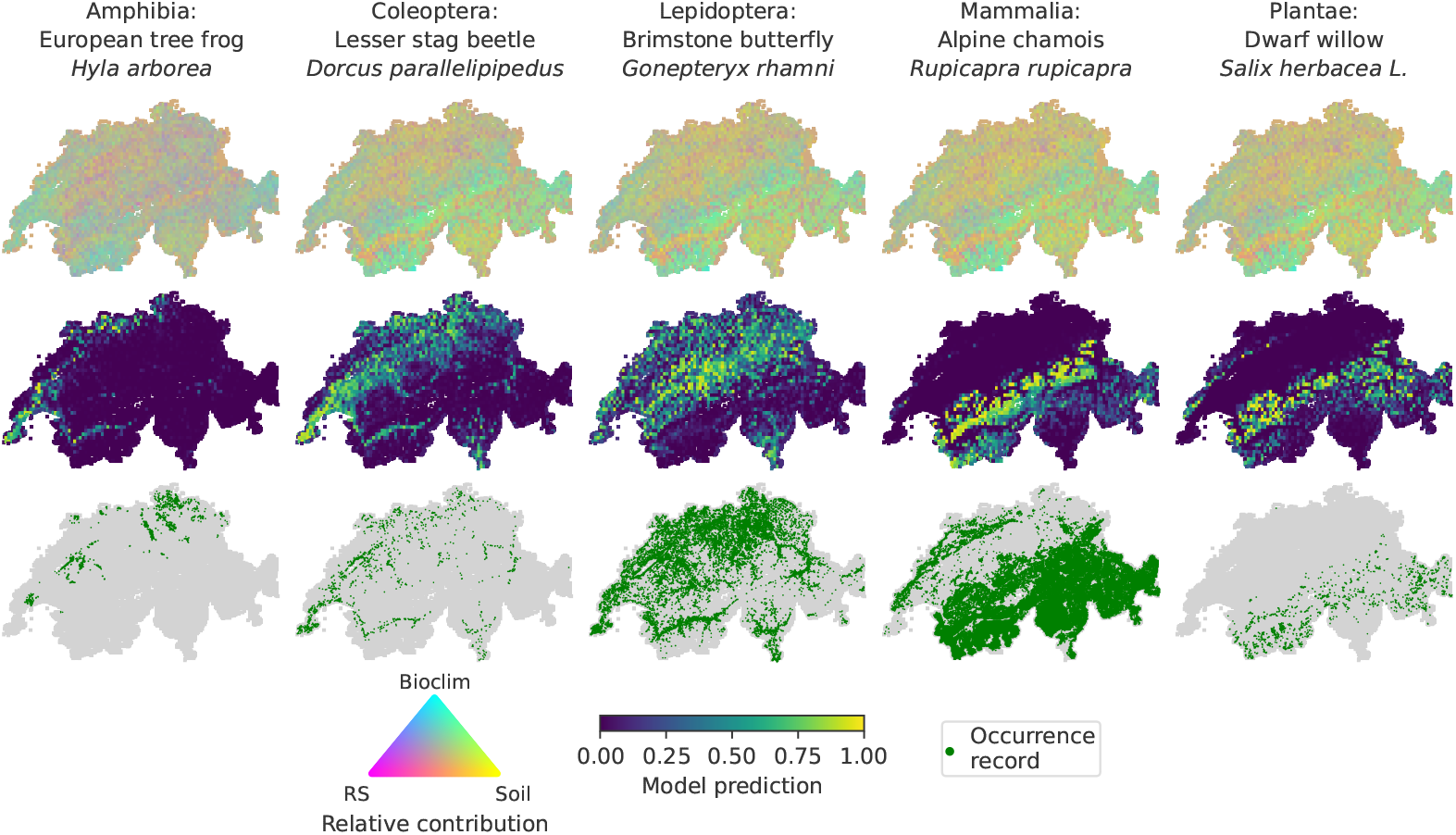
(a) Map of relative contribution of bioclimatic, soil, and remote sensing variables mapped to cyan, yellow, and magenta, respectively, (b) predicted distribution maps, and (c) species observation samples, for one example species from each of the taxonomic groups.

## 4. Discussion

In this study, we apply a CNN-based SDM framework to a Swiss dataset with high taxonomic diversity, consisting of both plant and animal species occurrence records. We investigate how scale affects model performance, comparing both single-scale and multi-scale modelling approaches for various predictor groups.

Our results show the strongest magnitude of scale effects and the largest gain from multi-scale approaches for remote sensing indices, while models trained on bioclimatic or edaphic variables and multimodal models have less pronounced scale effects. These results may be related to higher spatial autocorrelation in bioclimatic and soil variables compared to remote sensing indices and may also be attributed to the native resolution of these products. In the `SWECO25` dataset, remote sensing indices are resampled from 10 m resolution to 25 m, while the bioclimatic and soil variables are upsampled from 1 km and 100 m resolution, respectively. We find a strong similarity in relative scale effects across taxonomic groups for the various modalities. However, the absolute performance of single-scale models indicates that plant species prefer smaller scales, while animal species prefer larger scales. This is likely to be related to species mobility, with local predictors expected to be more meaningful for sessile organisms (Guisan and Thuiller, 2005). These results also reflect findings that high-resolution predictors improve model performance for plants Manzoor et al. (2018); Chauvier et al. (2022). However, in models using tabular data, increasing the resolution means decreasing the spatial extent. In contrast, using CNNs allows high-resolution predictors that capture the heterogeneous micro-environmental variations that are relevant to predicting plant distribution (Descombes et al., 2016) to be considered at larger spatial extents, capturing the spatial structure of the environment (Deneu et al., 2021).

Multi-scale models show superior performance to their single-scale counterparts across all taxonomic groups. Although the gain in absolute model performance is limited, our results also indicate that the performance of multi-scale models is more robust to individual scale selection, even when choosing an optimised single scale. In an approach analogous to ensemble modelling (Hao et al., 2019), the different spatial branches in our multi-scale models may learn complementary information. While the selection of the scales remains arbitrary, the inclusion of multiple scales may encourage the model to be more tolerant of non-informative scales. In many cases, the inclusion of multiple scales even leads to higher AUC scores than either of the corresponding single-scale models. These results reflect findings from a multi-scale SDM with tabular data in which focal statistics were used to vary the scale (Hallman and Robinson, 2020). Furthermore, they are consistent with the results from van Tiel et al. (2025), which used the same modelling framework on plant occurrence data across Europe from the GeoLifeCLEF 2023 dataset (Botella et al., 2023), albeit with drastically different predictor resolutions. The results from both studies indicate stronger effects of scale for remote sensing data, particularly in multi-scale settings, and a reduction in variability of model performance when including multiple scales.

Considering multiple modalities leads to better performance than unimodal models, suggesting that complementary information is learned from the separate modalities (Dollinger et al., 2024). There is a strong dependence of model performance on the choice of modalities. We find that scale effects are negligible in comparison to the effect of including the augmented set of modalities used in this study. While we limited the choice of modalities to three commonly used predictor types, many more predictors could be considered (Mod et al., 2016). Although covariates should be carefully selected to avoid overfitting, we acknowledge that some relevant modalities, such as topography, may have been left out here, and we expect their inclusion to lead to increases in model performance. Moreover, the flexibility of deep learning models enables the inclusion of various types of data in SDMs and similar multimodal approaches can easily be constructed combining spatial data with, for example, tabular data (Dollinger et al., 2024) or time series (Estopinan et al., 2022). Analogously to our study of spatial scale, the inclusion of time series would warrant a similar investigation of the effect of temporal scale.

Deep learning models have the potential of harnessing large amounts of species occurrence data, thanks to their scalability and ability to model arbitrarily complex relationships (Botella et al., 2018). These models also have the ability to model a large number of species with a single model (Cole et al., 2023; Botella et al., 2023). While most SDM studies using deep learning have focused on a single taxonomic group, here we successfully model species from different taxonomic groups with a single model. We note that model performance is higher for plants than for animals, although overall performance is found to be satisfactory. We suspect that the high similarity of contribution of the modalities and scales across taxonomic groups is primarily driven by plants, as they represent approximately 70% of the taxa considered here. The lack of variation in contributions indicates that the model learns that the same features are important for predictions across taxa, suggesting that the underlying complexity of biogeographic patterns is not captured. We leave to future work the investigation of the effect of modelling different species groups together or separately. Nonetheless, modelling many species across taxonomic groups together may allow perspectives towards integrating community-level information in a model with more ease than with single-species models (Ovaskainen et al., 2017).

Our results highlight the performance of multi-scale and multimodal neural networks. These complex models are more robust to individual scale or modality selection. Integrating multiple scales can drastically improve model performance, even for optimal single-scale selection. Furthermore, our use of gradient-based attribution methods reveals which predictors and scales contribute most to the model decision. Scale effects differ across predictor types, with the strongest effects found for remote sensing indices. The application of our method to a taxonomically diverse dataset suggests that smaller scales are more appropriate for sessile species, and mobile species benefit from larger scales. However, our robust multi-scale, multimodal models performed best across all taxonomic groups.

