## Supplementary material for "Multi-scale neural networks enhance species distribution modelling across predictors and taxonomic groups"

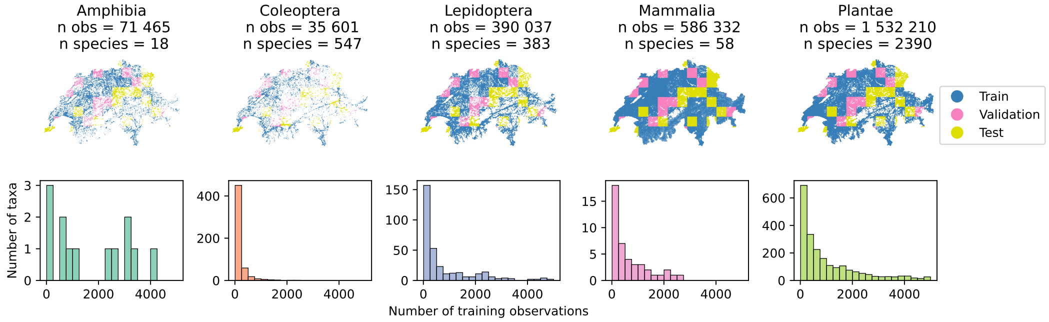


Figure S1: Occurrence data distribution per taxonomic group. (a) Geographic distribution of training, validation, and test splits with spatial blocks. (b) Histogram of the number of occurrences per species in the training set.

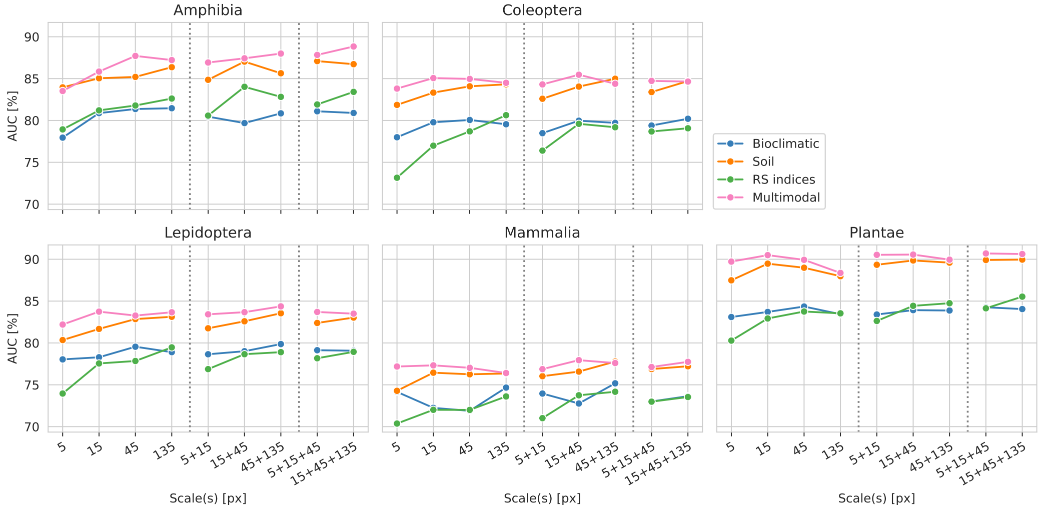


Figure S2: Median AUC scores (%) per taxonomic group evaluated on the test set for different scales for bioclimatic variables, soil variables, remote sensing (RS) indices, and all of the above (Multimodal). Dotted lines separate single scales, scale pairs, and triplets.

| Modality | Scale(s) | Overall | Amphibia | Coleoptera | Lepidoptera | Mammalia | Plantae |
|---|---|---|---|---|---|---|---|
| | 5 | 81.5 | 77.9 | 78.0 | 78.0 | 74.2 | 83.1 |
| | 15 | 82.3 | 80.9 | 79.8 | 78.3 | 72.2 | 83.7 |
| Bioclim | 45 | 82.6 | 81.4 | 80.1 | **79.5** | 71.9 | **84.4** |
| | 135 | 82.1 | **81.5** | 79.5 | 78.9 | **74.7** | 83.4 |
| | 5, 15 | 81.8 | 80.5 | 78.5 | 78.6 | 74.0 | 83.4 |
| | 15, 45 | 82.5 | 79.7 | 80.0 | 79.0 | 72.8 | 83.9 |
| | 45, 135 | 82.5 | 80.8 | 79.7 | 79.9 | 75.2 | 83.9 |
| | 5, 15, 45 | **82.7** | 81.1 | 79.4 | 79.1 | 73.0 | 84.3 |
| | 15, 45, 135 | 82.6 | 80.9 | **80.2** | 79.1 | 73.6 | 84.0 |
| | 5 | 85.8 | 84.0 | 81.9 | 80.3 | 74.3 | 87.5 |
| | 15 | 87.7 | 85.0 | 83.3 | 81.7 | 76.4 | 89.5 |
| Soil | 45 | 87.6 | 85.2 | 84.1 | 82.9 | 76.3 | 89.0 |
| | 135 | 86.6 | 86.4 | 84.3 | 83.1 | 76.4 | 88.0 |
| | 5, 15 | 87.4 | 84.9 | 82.6 | 81.7 | 76.0 | 89.3 |
| | 15, 45 | 88.3 | 87.0 | 84.0 | 82.6 | 76.6 | 89.9 |
| | 45, 135 | 88.4 | 85.6 | **85.0** | **83.5** | **77.8** | 89.6 |
| | 5, 15, 45 | 88.2 | **87.1** | 83.4 | 82.4 | 76.9 | 89.9 |
| | 15, 45, 135 | **88.5** | 86.7 | 84.7 | 83.0 | 77.2 | **90.0** |
| | 5 | 78.4 | 78.9 | 73.2 | 74.0 | 70.4 | 80.3 |
| | 15 | 81.3 | 81.2 | 77.0 | 77.6 | 72.0 | 82.9 |
| RS indices | 45 | 82.3 | 81.8 | 78.7 | 77.8 | 72.0 | 83.8 |
| | 135 | 82.4 | 82.6 | **80.6** | **79.5** | 73.6 | 83.5 |
| | 5, 15 | 81.2 | 80.6 | 76.4 | 76.9 | 71.0 | 82.6 |
| | 15, 45 | 83.0 | **84.0** | 79.6 | 78.7 | **73.7** | 84.4 |
| | 45, 135 | 83.2 | 82.8 | 79.2 | 78.9 | 74.2 | 84.7 |
| | 5, 15, 45 | 82.5 | 81.9 | 78.7 | 78.2 | 73.0 | 84.1 |
| | 15, 45, 135 | **83.9** | 83.4 | 79.1 | 78.9 | 73.5 | **85.5** |
| | 5 | 88.1 | 83.5 | 83.8 | 82.2 | 77.2 | 89.7 |
| | 15 | 89.0 | 85.8 | 85.1 | 83.7 | 77.3 | 90.5 |
| Multimodal: | 45 | 88.6 | 87.7 | 85.0 | 83.3 | 77.0 | 89.9 |
| Bioclim + Soil + RS | 135 | 87.3 | 87.2 | 84.5 | 83.7 | 76.4 | 88.4 |
| | 5, 15 | 88.9 | 86.9 | 84.3 | 83.4 | 76.9 | 90.5 |
| | 15, 45 | 89.1 | 87.4 | **85.5** | 83.7 | **77.9** | 90.6 |
| | 45, 135 | 88.5 | 88.0 | 84.4 | **84.4** | 77.6 | 89.9 |
| | 5, 15, 45 | 89.1 | 87.8 | 84.7 | 83.7 | 77.1 | **90.7** |
| | 15, 45, 135 | **89.2** | **88.8** | 84.6 | 83.5 | 77.7 | 90.6 |

Table 1: Median AUC scores (%), across all species and per taxonomic group, evaluated on test set for models considering different scales and modalities. Bold numbers indicate the best AUC per taxonomic group and per modality.
